# The p53-Mdm2 interaction through the lens of antithetic feedback

**DOI:** 10.64898/2026.10.04.756583

**Authors:** Wouter Jongeneel, Martina Scolamiero, Karl Henrik Johansson

## Abstract

We show that the p53-Mdm2 gene interaction can be understood as a form of antithetic feedback, but with mixed feedback loops. In particular, we show that this viewpoint sheds light on the emergence and robustness properties of p53’s oscillatory behaviour, as observed experimentally.

## I. Introduction

The p53 gene (a transcription factor) is involved in DNA repair and mutated p53 is linked to ≥ 50% of human cancers [1]. Oscillatory behaviour of p53 is understood to be linked to its functionality [2], [3], [4], however, the mechanism behind these oscillations is not fully understood. To elaborate, in normal conditions, the concentration of p53 is kept low (*i*.*e*., homeostasis), but at the onset of DNA damage (*e*.*g*., after irradiation), p53 starts to oscillate. What is more, the period of these oscillations is shown to be robust, whereas the amplitude can be highly variable. Again, it is not clear what leads to this form of robustness. A remarkable observation is that p53 oscillates in other species (*e*.*g*., in monkeys, dogs, mice and rats), may it be at a different frequency, but again with a robust period [5]. This has interesting evolutionary implications: the robustness of these oscillations should be a structural feature. Hence, we are interested in understanding what structural features of the underlying gene regulatory network (GRN) lead to these oscillations. Here, we are especially interested in network motifs [6].

The robustness of these types of oscillations has been discussed, *e*.*g*., in [7], however, not based on a physiologically meaningful model. The role and importance of positive feedback, to generate and sustain robust oscillations, is also an active topic of study in the realm of p53 [8], although often with the focus on stochastic simulations, *e*.*g*., see [9]. More recently, towards explaining robustness, a notion of buffering has been proposed [10]. An elementary explanation of the observed robustness is largely missing, nonetheless.

Now, although p53 is part of a large GRN, it is known that the E3 ubiquitin ligase Mdm2 is largely responsible for the regulation of p53 [11], [12]. Then, building upon [13], in this note we argue that the antithetic feedback motif [14], but with *mixed* feedback, fits the p53-Mdm2 interaction well. For instance, the transition from homeostasis to a sustained oscillation fits a Hopf bifurcation. Moreover, using this motif, an elementary explanation of the observed robustness can be obtained: p53 can be understood to act as an *actuator*.

### A. Contribution and structure

In this note we provide the following structural insights. We interpret the p53-Mdm2 interaction as antithetic feedback with mixed feedback. This network naturally accommodates homeostasis and oscillatory behaviour, plus, different roles of p53 and Mdm2 become apparent. In particular, we show that including complex forming (*e*.*g*., the binding of p53-Mdm2) is essential to make the above transparent. Moreover, through an interpretation as a control system and by an algebraic formula for the period of the limit cycle, robustness can be studied. In Sec. II we highlight the basics of integral control and in Sec. III we discuss basic properties of antithetic feedback. In Sec. IV we study the p53-Mdm2 interaction and provide our main results. We conclude in Sec. V.

## II. Integral control

In this section we illustrate a few well-known facts that aid in interpreting the p53-Mdm2 interaction later on. We point a reader unfamiliar with basic control concepts to [15].

First, recall that *integral* (I) controllers can oscillate, *e*.*g*., consider the most basic manifestation of an I controller for some gain *k*_*I*_ *>* 0:

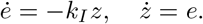

Here, “*e*” is understood as some *error* signal, with “*z*” its integral. The I controller is such that the error *e* follows 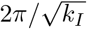-periodic dynamics, see Fig. 1 (A). Introducing a *proportional* (P) gain *k*_*P*_ *>* 0, that is,

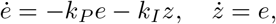

means the error dynamics are stabilized for any choice of *k*_*P*_, *k*_*I*_ *>* 0, albeit, with a oscillatory transient if 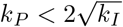, see Fig. 1 (B). Next, consider a *perturbed* control system of the form:

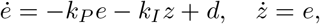

for some disturbance *d* ∈ ℝ. Due to the non-zero gain *k*_*I*_, the asymptotic dynamics are equivalent to the above. The difference is a change in setpoint, that is, the equilibrium is not (0, 0), but (0, *d/k*_*I*_), see Fig. 1(C). Next, consider the “*leaky*” I controller, as given by

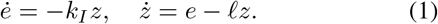

Such a system is important in the context of biology as every state exhibits some form of degradation/dilution. Despite stability, if we consider a perturbation *i*.*e*., *ė* = *− k*_*I*_ *z* + *d*, the equilibrium becomes ((*ld*)*/k*_*I*_, *d/k*_*I*_) and hence the leakiness breaks the integrator to the point that for *l →* +*∞* we have *e*\**→ + ∞*, see Fig. 1(D)-(E). Lastly, consider a leaky PI controller:

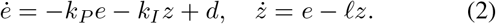

**Fig. 1:**
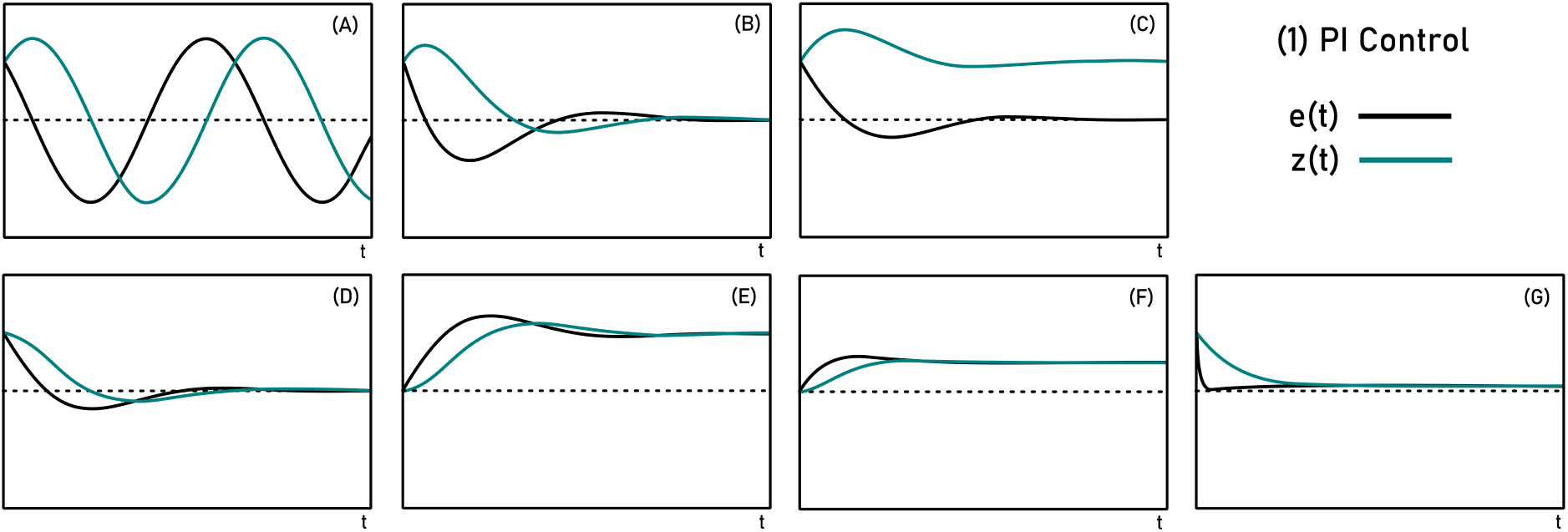
**(1)**: Examples related to Sec. II. (A): *k*_*I*_ = 1, (*e*(0), *z*(0)) = (1, 1), *t* ∈ [0, 10]. (B): (A) with *k*_*P*_ = 1. (C): (B) with *d* = 1. (D): (A) with *l* = 1. (E): (D) with *d* = 1 and (*e*(0), *z*(0)) = (0, 0). (F): (E) with *k*_*P*_ = 1. (G): (F), but with *k*_*P*_ = 10.

The equilibrium becomes (*ld/*(*lk*_*P*_ + *k*_*I*_), *d/*(*lk*_*P*_ + *k*_*I*_)). Hence, sufficiently high controller gains can overcome leakiness. Indeed, to overcome this form of leakiness, high-gain feedback has been proposed in the context of synthetic biology [16]. See Fig. 1(F)-(G) for examples. So far, we looked at *negative* feedback, when introducing *positive* feedback, the stabilizing effect of the leaks is important.

## III. Antithetic feedback

Antithetic feedback emerged as the biological manifestation of integral control [14]. Below we study the most basic version, that is, with just a single node representing the plant. It turns out that this structure is helpful in studying the p53-Mdm2 interaction. Indeed, it is known that *standard* antithetic feedback applied to a plant with 2 nodes may result in a sustained oscillation [14, Thm. S8.3]. The situation for just a single node is different, as discussed below. As we will focus on the p53-Mdm2 interaction, we also need not be concerned with stochastic models, as argued in [13, p. 6].

### A. Idealized antithetic feedback

First, we recall that the idealized antithetic feedback control ((I) in Fig. 2) of a plant with a single node is always stable. In [17, p. 52] such a result was shown “in the limit of strong binding”, that is, for large *µ*. See also [18].

**Fig. 2:**
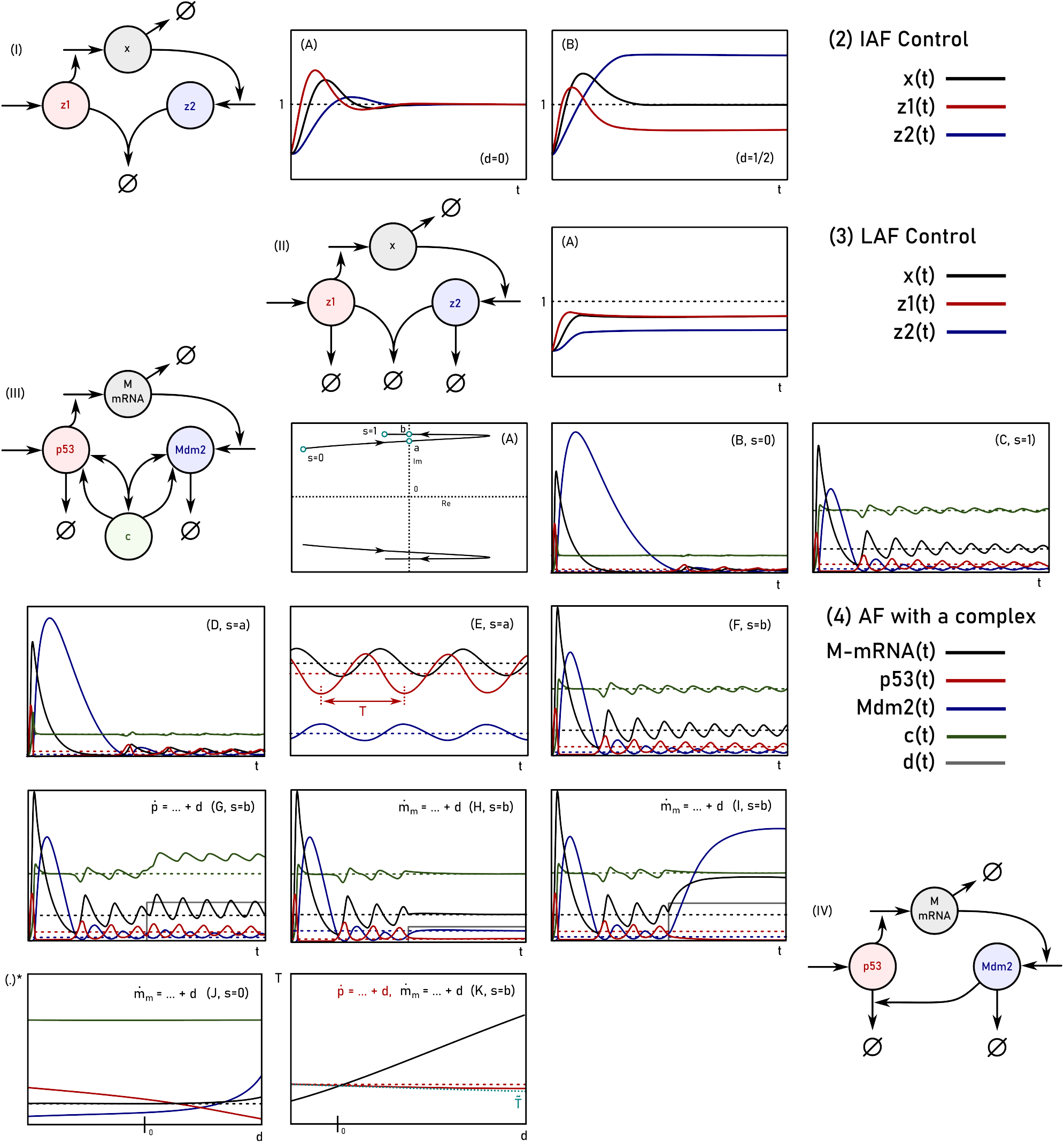
**(2)**: Examples related to Sec. III-A (*Idealized Antithetic Feedback* (IAF)). (A): *γ* = 1, *k* = 1, *r* = 1, *µ* = 1, *θ* = 1, (*x*(0), *z*_1_(0), *z*_2_(0)) = (0, 0, 0), *t* ∈ [0, 25] (see (3)). (B): (A) with *d* = 1*/*2. — **(3)**: Examples related to Sec. III-B (*Leaky Antithetic Feedback* (LAF)). (A): as in (2A) but with *l*_1_ = 1, *l*_2_ = 1 (see (4)). — **(4)**: Examples related to Sec. IV (the p53-Mdm2 interaction through *Antithetic Feedback with a complex*). We observe (supercritical) Hopf bifurcations when looking at (6) under [0, 1] ∋*s ↦ θ*(*s*) = (1 *− s*)*θ*_0_ + *sθ*_1_. (A): a pair of conjugate eigenvalues crosses the imaginary axis twice. (B-C): homeostasis. (D,F): a Hopf bifurcation. (E): when zooming in on (D) we see sustained oscillations (a stable limit cycle). (B-F) show trajectories starting from (*p*(0), *m*_*m*_(0), *c*(0), *m*(0)) = (1, 1, 1, 1), for *t* ∈ [0, 30], with (E) zooming in on the time-interval [85, 95] (*i*.*e*., after the transient). (G): the effect of a step disturbance at *t* = 15 (denoted by *d*) on 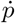 (p53). (H-I): the effect of a step disturbance on 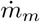 (Mdm2 mRNA). (J) equilibria as a function of *d ∈* [*−*10, 10]. (K): the period *T* as a function of *d ∈* [*−*10, 40], perturbing either 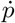 or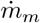. — Plots are made using MATLAB and Inkscape.

Let *d* ∈ ℝ_*≥*0_ be a constant and unknown disturbance, then, we study a dynamical system of the form

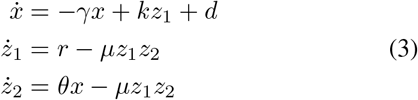

Here, “*x*” should be understood as the plant state, whereas “*z*_1_” and “*z*_2_” are the controller species (actuator and sensor, respectively). Compactly expressed, we denote the vector field (3) by *F*_1_. Then, see that *F*_1_ has a single equilibrium, being 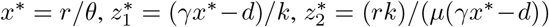.

Note in particular that *x*\* is independent of the disturbance *d* and the model parameters *γ* and *k*, hence the interpretation as integral feedback. Additionally, see that 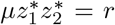, the feedback term *only* depends on *r*, asymptotically. Then, to make sure that *z*_1_ and *z*_2_ are not pathologically negative under large *d*, we have to assume that *d < γx*\* = (*γr*)*/θ*. Observe that there is a clear lack of symmetry in how the disturbance affects the controller species, *e*.*g*., *O*(*x*\* *− d*) versus *O*((*x*\* *− d*)^*−*1^). See Fig. 2 (2A-B) for an example.

Regarding stability, linearizing around 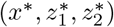 yields

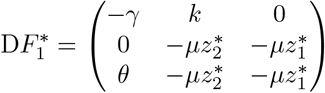

We can compute the characteristic polynomial of 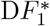, being *p*_1_(*s*) := *s*^3^ + *a*_2_*s*^2^ + *a*_1_*s* + *a*_0_, for 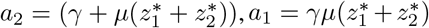 and 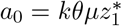. Then, a straight-forward computation and the Routh-Hurwitz stability criterion (*i*.*e*., *a*_2_*a*_1_ *− a*_0_ *>* 0) implies the following.

#### Proposition III.1

(Stability of ideal antithetic feedback). *Let d <* (*γr*)*/θ, then, the dynamical system* (3) *has a unique equilibrium point, being an element of* 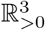. *This point is exponentially stable*.

#### Remark III.1

(On positive idealized antithetic feedback). Note that for the system (3), if we pick *µ <* 0 instead of *µ >* 0 then 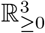 will contain no equilibrium point.

To continue, understanding the *global* dynamics of (3) turns out to be hard. Numerical studies are provided in [18]. By Proposition III.1, we know that Hopf bifurcations cannot occur, and this indicates that (3) is not a plausible model for the p53-Mdm2 interaction. Nevertheless, (3) may still display limit cycles (*e*.*g*., as curl(*F*_1_) ≠ 0, *F*_1_ is no gradient flow). However, despite some particular comments one can make via [19], [20], characterizing the *global* dynamics is open.

### B. Leaky antithetic feedback

The above is *idealized*, the controller species do not degrade. Next, we consider a “leaky” antithetic feedback controller ((II) in Fig. 2). As with PI control, the setpoint changes, the stability does not, *cf*. (1). This setting is particularly relevant when considering *positive* antithetic feedback (*i*.*e*., the leaks aid the stabilization). To that end, consider

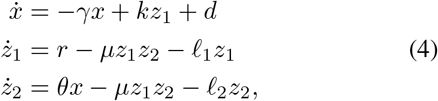

for some *l*_1_, *l*_2_ *>* 0. One observes that we have again a unique equilibrium in 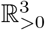, plus forward invariance in 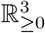.

In particular, see that as *ż*_1_ *− ż*_2_ = *r − θx −* (*l*_1_*z*_1_ *− l*_2_*z*_2_) we get that 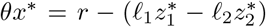. There is a potential offset in regulated setpoint due to the leakiness. See Fig. 2 (3A) for an example. Regarding the equilibrium, we get

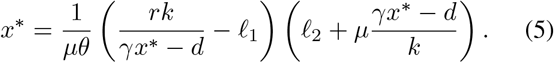

Now rewrite (5) into an equation quadratic in 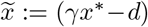:

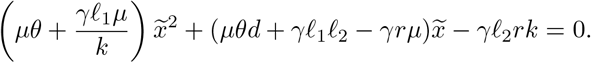

Then, recover *x*\* through the positive root. When *l*_2_ *>* 0, such a root always exists and is unique. When *l*_2_ = 0, *d*≤ (*γr*)*/θ* guarantees again the existence of such a solution. Next, we get 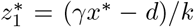, which is always positive. On the other hand 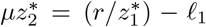. Comparing to the idealized case, where 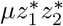 was equal to *r*, we see that in the leaky case, the feedback term is not readily invariant under a change in parameters. To continue, to see that 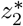 is also positive, we use the relation 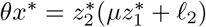.

Regarding stability, linearizing around 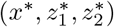 yields

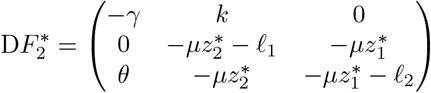

We can compute the characteristic polynomial of 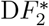, being *s* ↦ *p*_2_(*s*) := *s*^3^ + *a*_2_*s*^2^ + *a*_1_*s* + *a*_0_, for 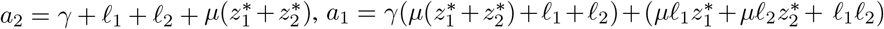 and 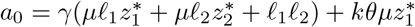.

Similar to the previous section, one may follow again a Routh-Hurwitz argument to conclude on stability. To that end, let 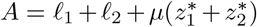 and 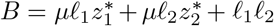. Then, observe that

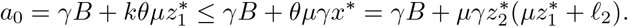

Now, as *a*_2_*a*_1_ = (*γ* +*A*)(*γA*+*B*) = *γ*^2^*A*+*γA*^2^ +*AB* +*γB*, we simply observe that *γA*^2^ + *γB ≥ a*_0_ and *γ*^2^*A* + *AB >* 0. Hence *a*_2_*a*_1_ *− a*_0_ *>* 0 and we have shown the following.

#### Proposition III.2

(Stability of leaky antithetic feedback). *Let d <* (*γr*)*/θ, or, when d* ≥ 0, *let l*_2_ *>* 0, *then, the dynamical system* (4) *has a unique equilibrium point, being an element Of* 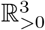. *This point is exponentially stable*.

#### Remark III.2

(On leaky positive antithetic feedback). Without leaks, a positive feedback does not result in a feasible equilibrium point, *i*.*e*., equilibria fail to be elements of 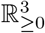. However, with sufficiently large leaks, we again have a feasible set of equilibrium points.

### C. Antithetic feedback with a complex

Instead of models like (3), one may include the complex formed by *z*_1_ and *z*_2_, doing so leads to models like the one shown in Fig. 2(III). It is this level of extra information that aids us in studying the p53-Mdm2 interaction. An important aspect of such a model is that now we have a form of *mixed* feedback (*i*.*e*., both negative- and positive feedback loops).

Mixed feedback systems received a lot of attention, especially towards understanding oscillations [21]. Here, it is especially interesting to study how qualitative behaviour can change, without changing the “anatomy” of the system [22]. Indeed, we aim to understand how to transition from homeostasis to a sustained oscillation, without altering the network. What is more, based on the precise type of nonlinearity, mixed feedback systems are typically studied as Lur’e systems, in particular by means of dominance theory [23].

In the next section, we study the p53-Mdm2 specific case.

## IV. The p53-Mdm2 interaction

In this section we start by discussing the rationale behind studying the p53-Mdm2 interaction via an antithetic feedback motif. After doing so, we study mathematical ramifications.

To start, we provide several comments on experimentally observed behaviour and modelling choices.

### (i) Focus on Mdm2

Mdm2 is known to control p53 by binding to it (mono- or poly ubiquitination) [12]. Experiments with mice confirmed that Mdm2 is essential here [11]. As p53 is known to be part of a larger network, our aim is to understand a “minimal network” first.

### (ii) Sustained oscillations

Experiments show that p53 oscillates after irradiation. These oscillations appear to be *damped* when averaging over a population [2]. However, looking at individual cells, sustained oscillations are observed, with a robust period and variable amplitude [3].

### (iii) Delays

Several works contain models with a delay. However, delays obfuscate how structural features of the GRN introduce oscillations, *e*.*g*., the single-node network described by 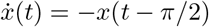 oscillates.

### (iv) Noise, variability and robustness

Although the period is seemingly robust, the nominal period varies between species [5]. The authors in that paper also mention that “*It is possible that the elements under selection are not oscillations per se, but rather strong feedback controls*”.

### (v) Saturation

Other works study a simplified and saturated (*e*.*g*., to induce delays) version of the model that we study, *e*.*g*., see [24], [25] and Fig. 2(IV). They assume that the p53-Mdm2 complex is already in equilibrium, this, however, obfuscates the integral feedback structure.

With the above in mind, we follow [13]. Their model can be depicted as Fig. 2(III) (although it was depicted differently in the original paper *cf*. [13, Fig. 1]; we emphasize the connection to antithetic feedback). Here, the states correspond to concentrations of: nuclear p53 (*p*), Mdm2 mRNA (*m*_*m*_), Mdm2 (*m*) and the p53-Mdm2 complex (*c*).

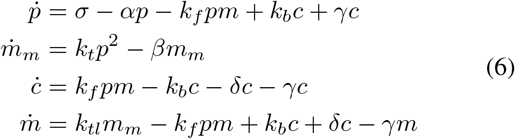

For the full motivation of (6) we point the reader to [13], but we mention that *σ* is the constant production rate of p53 (with the assumption being justified by neglected interactions being slow), *k*_*t*_ is the transcription rate, *k*_*tl*_ the translation rate, *β* the degradation rate of the Mdm2 mRNA, *k*_*f*_ is the complex (p53-Mdm2) formation rate whereas *k*_*b*_ is the complex break-up rate, *δ* is the Mdm2-mediated degradation rate of p53, with *α* containing all other forms of p53 degradation. On the other hand, Mdm2 degrades at a rate *γ*, irrespective of being bound to p53.

Comparing (6) to earlier sections on antithetic feedback, see that we have the following relation of equilibria

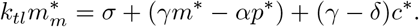

That is, we see that the regulation of 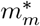 is altered by the leakiness of the antithetic feedback (*i*.*e*., (*γm*\* *− αp*\*)), plus the positive feedback from the complex (*i*.*e*., (*γ − δ*)*c*\*).

#### DNA damage

Following [13], after irradiation, DNA damage, leads to increased auto ubiquitination of Mdm2, decreased ubiquitination of p53 by Mdm2 and a weaker binding of p53 and Mdm2. In the context of (6), this means an increase of *γ*, a decrease of *δ* and an increase of *k*_*D*_ = *k*_*b*_*/k*_*f*_ (decrease of *k*_*f*_). One may add that there is experimental evidence that *k*_*t*_ goes down under hypoxia.

### A. Properties of (6)

First, see that 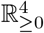 is invariant under (6), we will call the corresponding vector field *F*_0_, and this vector field is parametrized by 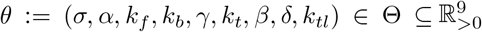, for Θ being some open subset. Then, in [13, p. 2], the authors state that their model displays oscillations because of the single negative feedback loop. We would like to sharpen this statement, the oscillations occur because of the *mixed* feedback. This is particularly transparent when studied through the lens of *antithetic feedback*.

#### Lemma IV.1

(Existence, uniqueness and smoothness of equilibria). *For any choice of θ* ∈ Θ, *the dynamical system* (6) *has a unique equilibrium point, denoted* 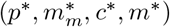. *This point is an element of* 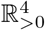. *Moreover*, 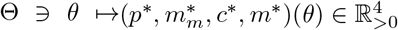 *is C*^*ω*^*-smooth*.

*Proof*. To study equilibria, we observe from (6) that at least the following relations must hold:

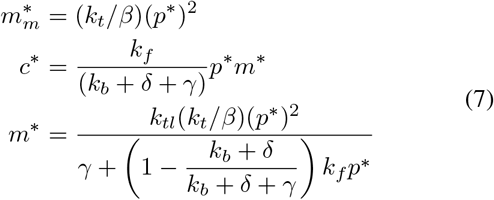

For *p*\*, we get from (6) and the above that

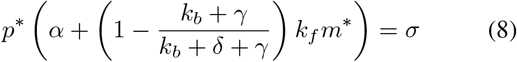

Then, a straight-forward application of Descartes’ law of signs [26, Thm. 2.33] shows that there is a unique positive *p*\* that complies with (6), *i*.*e*., one transforms (8) to a cubic polynomial in *p*\*, *e*.*g*., *p*\* ↦*a*_3_(*p*\*)^3^ + *a*_2_(*p*\*)^2^ + *a*_1_*p*\* + *a*_0_, with signs (+, +, *a*_1_, *−*), so regardless of the precise value of *a*_1_, there is always one sign change and hence one strictly positive root. In its turn, this implies that, for any *θ ∈* Θ, (6) has a unique equilibrium point 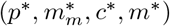in 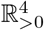. What is more, since by the above, *p*\* corresponds to a *simple* strictly positive root, *θ* ↦*p*\*(*θ*) is *C*^*ω*^ by the real-analytic implicit function theorem [27, Thm. 1.8.3]. Hence 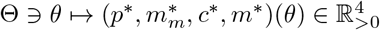 is *C*^*ω*^.

Lemma IV.1 already limits bifurcations complying with (6), simplifying a study of oscillations. Next, we characterize stability of the equilibrium. In contrast to the models studied above, instability is possible, hence, generating sustained oscillations is not readily obstructed. Note, we did not introduce delays, oscillatory driving inputs or a saturation.

Then, the following lemma is stating that the equilibrium is either stable, or unstable with an oscillatory component. Below, we will exploit the coefficients of the polynomial.

#### Lemma IV.2

(Stability of equilibria). *The characteristic polynomial corresponding to the strictly positive equilibrium point under* (6) *has either* 0, 2, *or* 4 *real roots that are strictly negative, but no real roots that are strictly positive*.

*Proof*. To start, the linearization of *F*_0_ around any (equilibrium) point is 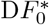 and given by the matrix

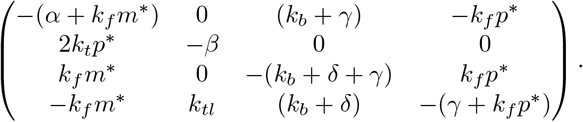

The sign structure of 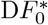 is usually exploited to study oscillations, however, as node *m* is of degree 3, we cannot readily conclude on 2-cooperativity *cf*. [28, Sec. III].

To study local stability, we compute det 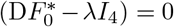. It is convenient to explicitly write *λ* ↦ Det 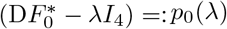 where *p*_0_(*λ*) =: *λ*^4^ + *b*_3_*λ*^3^ + *b*_2_*λ*^2^ + *b*_1_*λ* + *b*_0_ for

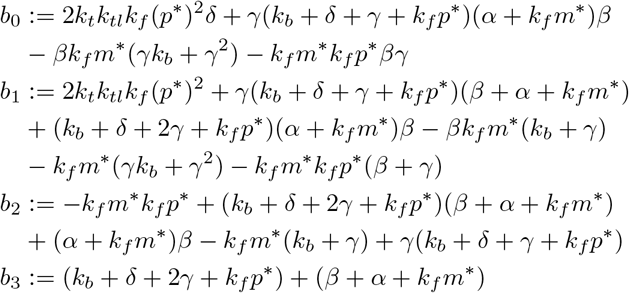

One can check that all coefficients of *p*_0_ are strictly positive, for any *θ* ∈ Θ. Moreover, Descartes’ law of signs [26, Thm. 2.33] implies we cannot have strictly positive real roots, that is, unstable modes must have oscillatory components. Now, applying Descartes’ law to *λ* ↦*p*_0_(*−λ*) implies that we have either 0, 2, or 4 real roots that are strictly negative.

Lemma IV.2 indicates that one should expect a Hopf bifurcation, *e*.*g*., see [29, Sec. 3]. For example, in [13], the choice of parameters (recall, *θ* = (*σ, α, k*_*f*_, *k*_*b*_, *γ, k*_*t*_, *β, δ, k*_*tl*_))

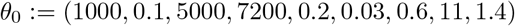

is chosen to represent homeostasis, whereas the selection

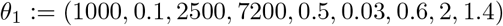

should correspond to the behaviour after DNA damage. Indeed, introduce the parameter *s* ∈ [0, 1] and study (6) along [0, 1] ∋*s* ↦*θ*(*s*) := (1 *−s*)*θ*_0_ + *sθ*_1_, this reveals the following: one crosses the imaginary axis twice. The first Hopf bifurcation occurs at *s ≈* 0.262 (point *a*), with *ω* ≈ 1.88 (*i*.*e*., the crossing occurs at *±iω*), leading to oscillations with a period *T ≈* 3.34 (*i*.*e*., *T* = 2*π/ω*), as shown in Fig. 2 (4) (A,D,E). The second bifurcation occurs at *s ≈* 0.973 (point *b*), with *ω ≈* 2.09, see Fig. 2 (4) (A,F).

To gain more insights in how *ω* depends on *θ*, we can compute det 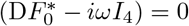. In particular, we must have *−ib*_3_*ω*^3^ + *ib*_1_*ω* = 0 and thus 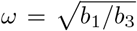. This algebraic expression provides insights, locally, in how the period of emerging periodic orbits depends on the parameter *θ*.

### B. On homeostasis

As (6) can be understood as mixed antithetic feedback, one might expect a form of robust perfection adaption (RPA) with regards to the *m*_*m*_ variable, at least for some choice of *θ* ∈ Θ. This does not directly follow from local stability.

One way to study this, without appealing to precise equations, is via the framework as developed by Golubitsy, Stewart and co-workers, *e*.*g*., see [30]. Suppose we have a smooth vector field on ℝ^*n*^ of the form:

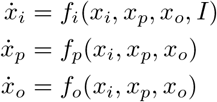

where “*i*” denotes a dedicated input node and “*o*” some output node. The rest is internal. In compact form we would write 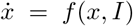. Suppose now there is an equilibrium point (*x*\*, *I*^*′*^) that is exponentially stable under the vector field *f*. In this case, 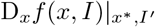 must be invertible (*i*.*e*., as the matrix is Hurwitz). Hence, by the Implicit Function Theorem (IFT) we have that there exist smooth functions *x*_*i*_(*I*), *x*_*p*_(*I*), *x*_*o*_(*I*) such that *f* (*x*(*I*), *I*) = 0 for all *I* in a neighbourhood of *I*^*′*^, ith *x*(*I*^*′*^) = *x*\*. Then, implicit differentiation yields that

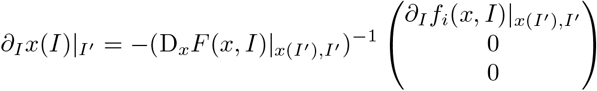

Rewriting and projecting onto *x*_*o*_ leads to

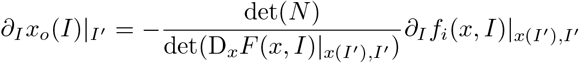

Where

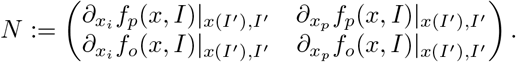

Then, we speak of *infinitesimal* homeostasis, for *x*_*o*_ under *I*, when 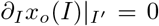. Studying this notion of homeostasis is of interest since some network structure appears and one can conclude independently of the precise choice of *f, i*.*e*., if 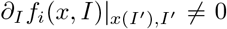, a necessary and sufficient condition for infinitesimal homeostasis becomes det(*N*) = 0.

For example, if we look at ideal antithetic feedback (3) and identify *d* with *I*, then, if we let *x* be both the input and the output node, we extract the appropriate matrix *N* and obtain det 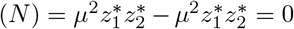. Indeed, we do have RPA there. Now, consider the p53-Mdm2 interaction (6). One readily checks that under additive perturbations entering through *m*_*m*_ (*i*.*e*.,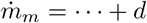), infinitesimal homeostasis does not occur for any set of parameters as

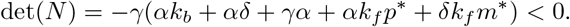

Nevertheless, if we plot how the equilibria move as a function of *d*, for example, for *θ*_0_, we see that the 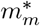 - curve is almost flat, see Fig. 2 (4) (J). We can speak of “*near-perfect homeostasis*” (see [30, Def. 2.1]), complying with what one expects from the antithetic framework.

When looking at the p53-Mdm2 interaction, a similar observation can be made. In Fig. 2 (4) (K), we compare what happens to the period *T*, around the Hopf bifurcation *θ*(*b*), under either 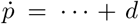 or 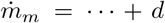, for a range of perturbations *d*. We observe a noticeable difference; perturbi<u>ng p5</u>3 has little effect on *T*. As *T* = 2*π/ω* and 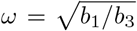, we can be more precise and study *∂*_*d*_*T* |_*d*=0_, that is, we can study infinitesimal period homeostasis. In general, one may derive an integral expression (*e*.*g*., see [30, §4.4]), however, more simply, it follows from Lemma IV.1 that if we consider 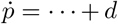, then both *∂*_*d*_*p*\* |_*d*=0_ *>* 0 and *∂*_*d*_*m*\* |_*d*=0_ *>* 0, and in fact, *∂*_*d*_*T* |_*d*=0_ *<* 0. So again, we need to speak of some form of *near-perfect* homeostasis.

### C. On robustness

Sec. IV-B showed that robustness is not readily captured by infinitesimal conditions. Hence, to get insights, we present some approximations.

Here, it is worthwhile to note that *θ* is not arbitrary. Following [13], we note that the set of parameters is restricted. In particular, we have *σ, k*_*f*_, *k*_*b*_ *≈* 10^3^, *δ ≈* 10^1^, *k*_*tl*_ *≈* 10^0^, *α, β, γ ≈* 10^*−*1^, *k*_*t*_ *≈* 10^*−*2^. This knowledge allows for an approximate robustness study of *ω*.

Given Fig. 2 (4) (K), and that experimental results indicate that p53 oscillates with a robust period but variable amplitude (after DNA damage), we study perturbations of the form 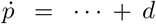. Now, it is convenient to interpret these perturbations as a perturbation to *σ, cf*. (6). See that for *σ ≪ α, p*\**<u>≈</u> σ/ α*. On the other hand, for *σ ≫ α* we have 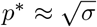. What is more, from (7) and (8) we get that *m*\* *≈* 10*k*_*t*_(*p*\*)^2^*/*(10^*−*1^ + *p*\*) and *p*\*(10^*−*1^ + *m*\*) *≈ σ*.

Hence,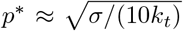. Then, 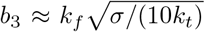, *b*_1_ *≈* 10^*−*1^*k*_*f*_ *σ* and thus 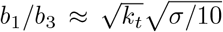. To illustrate this approximation, in Fig. 2 (4) (K), we show 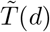 as computed via 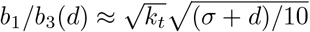.

Albeit coarse, this analysis points to why the period might be robust (*e*.*g*., compare 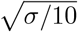 with 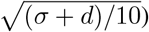. See Fig. 2 (4G) for a simulation result. This robustness also seems to align with the experimental observation that the amount of DNA damage does not translate to qualitatively different oscillations (although the number of pulses might be affected), *e*.*g*., see [31, Fig. 3]. What is more, from an operational viewpoint, the robustness might be explained through p53 acting as an “actuator”, with the perturbations relating to a change in setpoint, affecting the amplitude, but not the transient oscillatory behaviour. Looking at p53 through the lens of antithetic feedback enabled this view.

Regarding Mdm2 and disturbances of the form 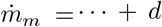, the antithetic feedback interpretation and near-perfect homeostasis indicate that a modest disturbance can be rejected. Yet, for a sufficiently large disturbance, we observe divergence from the setpoint, see Fig. 2 (4) (H-I). Observe that in both cases the oscillations vanish, which aligns with the physiological interpretation of the p53-Mdm2 interaction. Moreover, the dependence on *k*_*t*_, which should be understood as the *k*_*I*_ gain in an I-controller, appears naturally *cf*.

Sec. II. Indeed, in a standard I-controller, the *k*_*I*_ gain leads to oscillations with a frequency pr<u>op</u>ortional to 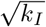. Here, the frequency is proportional to 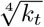 (due to the *p*^2^ term in (6)). This also seems to fit the hypoxia case from [13].

## V. Discussion and conclusion

In this note we have illustrated how the view from antithetic feedback may help in understanding the dynamical phenomena as observed in the context of the p53-Mdm2 interaction. Next, one should enlarge the network, *e*.*g*., by introducing Wip1 and ATM. From a theoretical viewpoint, we need new tools to capture the forms of *near*-perfect homeostasis as encountered in the above. Our current tools are too rigid, even for a model as simple as the one we consider in this work. What is more, we need to make the notion of “robustness” more precise and study if the oscillations are to be understood as truly sustained (compared to transient). Lastly, it is interesting to study these motifs in the light of recent generative results, *e*.*g*., see [32, Fig. 6], entrainment, buffering and other notions of controlled p53, also experimentally, *e*.*g*., see [10], [25].

